# The molecular landscape of polycystic kidneys is marked by common alterations in purine metabolism

**DOI:** 10.1101/2025.08.07.668867

**Authors:** Jean-Paul Decuypere, Daniel M Borras, Priyanka Koshy, Ludwig Missiaen, Steffen Fieuws, Nikky Corthout, Humbert De Smedt, Diethard Monbaliu, Jacques Pirenne, Tania Roskams, Bart Ghesquière, Bert Bammens, Abhishek D Garg, Djalila Mekahli, Rudi Vennekens

## Abstract

Autosomal dominant polycystic kidney disease (ADPKD) is the most common inherited kidney disease. Cysts develop through dedifferentiation of tubular epithelial cells, but the sequence of molecular events and their relative importance remain unclear. To address this gap in knowledge, 40 cysts from 4 ADPKD kidneys and 4 microcystic tissues were mapped on transcriptomic and histological level. Cyst were heterogenous and we identified 6 cystic subclusters with 2 deviations from the main trajectory, dependent on the rate of interstitial remodeling, inflammation and dedifferentiation. Loss of proximal tubular marker gene expression was more pronounced compared to those of other tubular segments. Altered expression of metabolic pathways was consistent among the cysts, which was further analyzed in human and mouse cell lines. Purine metabolism was similarly altered in all ADPKD cell lines, and its modulation with azathioprine suppressed cyst formation in vitro. In conclusion, by focusing on common altered pathways in cysts and cell models, we have identified purine metabolism as a novel potential target in ADPKD.

## INTRODUCTION

Autosomal dominant polycystic kidney disease (ADPKD) is the most common inherited kidney disease (Torres *et al*, 2007), with a prevalence of 1 in ca. 1000 individuals and accounting for 10% of the European dialysis population. It is most frequently (75-80%) caused by mutations in the *PKD1* gene encoding polycystin-1 (PC1) (Cornec-Le Gall et al., 2019; Hughes et al., 1995). Besides systemic manifestations such as liver cysts, hypertension and cardiac abnormalities, the most prominent phenotype is the gradual development of kidney cysts, concomitant with kidney enlargement and a decline in kidney function. This leads to kidney failure at an average age of ca. 50 years. The cysts develop due to dedifferentiation and enhanced proliferation of mutated tubular epithelial cells in all segments of the nephron. cAMP-driven fluid secretion into the lumen further promotes cyst swelling. This process is molecularly regulated by changes in Ca^2+^, cAMP, PKA and PI3K-Akt-mTOR signaling, among others (Torres & Harris, 2014). In addition, kidney cysts are characterized by ciliary dysfunction, loss of polarity, apoptosis, mitochondrial abnormalities, metabolic reprogramming, inflammation, oxidative stress and epigenetic changes (Chang & C M Ong, 2018; Ong & Harris, 2015; Podrini et al., 2018; Rowe et al., 2013; Song et al., 2009; Zimmerman et al., 2020). Despite the occurrence of all these alterations in cystic tissue or cells, the exact sequence of cellular and molecular events as well as their relative importance in the process of cyst formation remain unclear. Based on a transcriptomic and histological analysis of 40 individual cysts of 4 *PKD1* patients, we present a distinct heterogeneity in individual cysts, even within a single patient. In describing this heterogeneity, we determined the molecular and cellular events that occurred consistently in all cysts, or at different rates in the progression of ADPKD. In this manner, we identified and validated purine metabolism as a novel common and early pathway important in cystogenesis.

## RESULTS

### Differences between healthy tissue and ADPKD cystic membranes

We performed transcriptomic RNA-Seq analysis on 40 ADPKD cystic membrane samples from 4 *PKD1* patients (Suppl. Table 1), microcystic (MCT) samples from 2 *PKD1* patients, and 4 healthy kidney tissues (Suppl. Table 2). The cyst, MCT and healthy samples clearly separated from each other in the principal component analysis (Fig. 1A). In the cyst samples, 1139 genes were significantly down- and 1444 upregulated compared to the “Healthy + MCT” group. As expected, PKD1 was among the top downregulated upstream regulators (Fig. 1B). Other differentially regulated upstream regulators, such as AGT (angiotensin), TNF, IL1, IL6, NFκB and TGFB1 (Fig. 1C), have been described in ADPKD as well (Hian et al., 2016; Li et al., 2021; Mun & Park, 2016; Zimmerman et al., 2020). However, cystic tissues were histologically very different from healthy kidney tissue (Fig. 1D). Cysts were lined by a single layer of flattened epithelial cells surrounded by thickened interstitium consisting mostly of fibroblasts and collagen fibers. In addition, this thick interstitial layer contained inflammatory cells, capillaries, arteries, arterioles, adipocytes, atrophied tubuli or microcysts (Suppl. Fig. 1). Normal renal parenchyma, tubuli or glomeruli were rarely observed in these cyst specimens. As such, the strong transcriptomic differences between healthy *versus* cystic tissue likely primarily reflect the differences in histological constitution, which might obscure the changes crucial for cyst development.

**Fig. 1:**
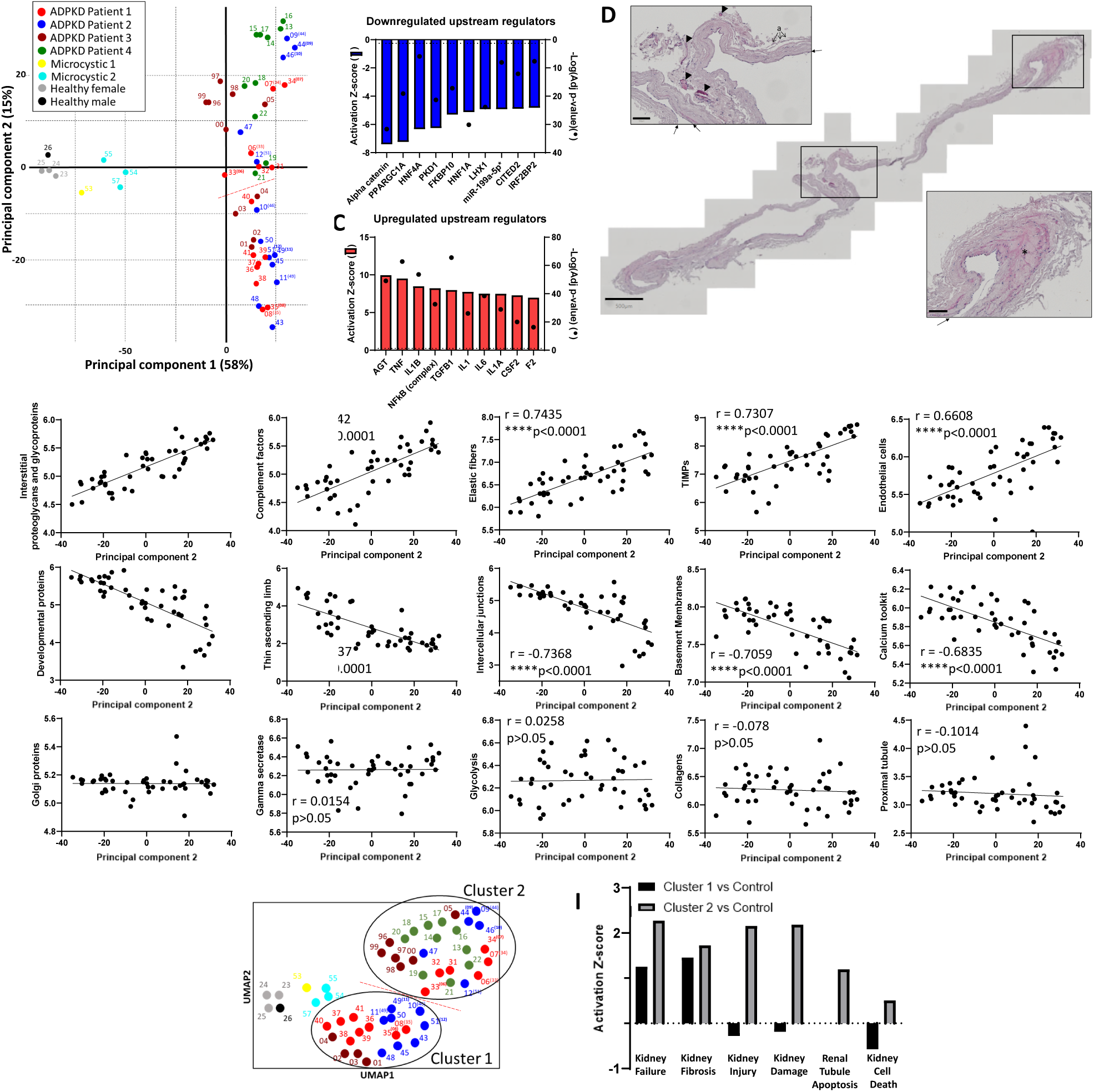
Heterogeneity among cyst samples from ADPKD patients. A) Principal component analysis of all analyzed samples. The red dashed line indicates the separation of the two clusters defined in the Leiden cluster analysis of panel H. All samples are assigned by a number, while the color corresponds to specific patients. For adjacent samples from the same cyst, the number of the other sample is indicated in brackets. B-C) The top 10 downregulated (B) and upregulated (C) upstream gene regulators according to pathway analysis on the cystic versus healthy + MCT samples. D) Example of an H&E staining of a cystic membrane. Arrows indicate the cyst-lining cells, triangles indicate capillaries and the asterisk marks fibrotic tissue. Scale bars are 500 µm (whole tissue) and 100 µm (insets). E-G) Correlation analyses of the values of principal component 2 with the top 5 significant molecular and cellular signatures, with indication of the Pearson’s r-value and p-value. Examples of significantly positive (E) and negative (F) correlations, or no correlations (G). H) Leiden clustering analysis revealed 2 clusters among the cyst samples. The red dashed line indicates the separation of the two clusters. All samples are assigned by a number. For adjacent samples from the same cyst, the number of the other sample is indicated in brackets. I) Pathway analysis comparison of significantly altered nephrotoxicological functions of cluster 1 and of cluster 2 versus healthy samples.

### Transcriptomic heterogeneity in cyst samples correlates with the ADPKD phenotype

We focused subsequently on the differences along the second principal component, explaining the heterogeneity between the cyst samples (Fig. 1A). The observed heterogeneity was independent of patient, cyst localization (cortical/medullary), size and fluid color (Suppl. Fig. 2). Although adjacent pieces from the same cyst mostly adjoined on the principal component plot (samples 44 and 09; samples 34 and 07; samples 33 and 06; samples 49 and 11; samples 35 and 08), this was not the case for 2 cysts (samples 51 and 12; samples 46 and 10), suggesting that the heterogeneity existed even within a single cyst. We performed a correlation analysis on the second principal component values for the cyst samples with various transcriptomic signatures relevant to ADPKD (Suppl. Table 3). A positive correlation existed with signatures related to the interstitium (e.g. interstitial proteoglycans and glycoproteins, elastic fibers, tissue inhibitors of metalloproteinases (TIMPs), endothelial cells, fibroblasts) and inflammation-related responses (e.g. complement system, apoptosis) (Fig. 1E and Suppl. Table 4), while strong negative correlations were related to kidney differentiation (developmental proteins, thin-ascending-limb markers, intercellular junctions, basement-membrane markers) and Ca^2+^ signaling (Fig. 1F). Collagens and macrophages (interstitium), and proximal or distal convoluted tubular markers (differentiation) did not significantly correlate with the heterogeneity, as well as expression of gamma-secretase members (which cleave PC1), metabolic pathways (e.g. glycolysis, fatty-acid oxidation, purine metabolism), signaling pathways (e.g. PI3K-Akt-mTOR, MAPK) and subcellular compartments (Golgi apparatus, microtubules, actin, nucleus, ER) (Fig. 1G and Suppl. Table 4). It should be noted that, although they did not differ between the different cystic samples, some of these hallmarks were significantly different compared to healthy tissue (see further).

To further analyze the significance of the transcriptomic heterogeneity between cysts, we performed a Leiden clustering analysis, revealing 2 distinct clusters among the cyst samples (Fig. 1H). The separation of the clusters on the UMAP plot (red dashed line in Fig. 1H) was also observed in the principal component plot (red dashed line in Fig. 1A). Comparative pathway analysis showed that the alterations in cluster 2 versus healthy tissue were more strongly associated with kidney failure, injury, fibrosis and cell death compared to cluster 1 (Fig. 1I).

### Main trajectory and deviations of cystic subclusters

To analyze the heterogeneity in more depth, we performed Single-cell Trajectories Reconstruction, Exploration And Mapping (STREAM) analysis (Sprooten et al., 2024). Among the cyst samples, 6 subclusters diverged (C1-C6) (Fig. 2A). The 2 larger clusters from the Leiden cluster analysis (Fig. 1H) were also apparent along the trajectory between C3 and C4 (red dashed line in Fig. 2A). Two deviations were apparent from the main trajectory: a deviation of C2 and between C5 and C6. Considering the significant correlations of the heterogeneity with interstitium and (de)differentiation (Fig. 1), we first examined the expression of general interstitial and tubular markers. Interstitial markers showed a gradual increase and tubular markers a gradual decrease over the pseudotime, with C5 having significantly higher tubular- and lower interstitial-marker expression versus C6 (Fig. 2B). Statistical significances of subcluster 2 versus subcluster 3 and of subcluster 5 versus subcluster 6 are shown in the graphs. The statistical comparison of all subclusters can be found in supplementary Table 5.

**Fig. 2.**
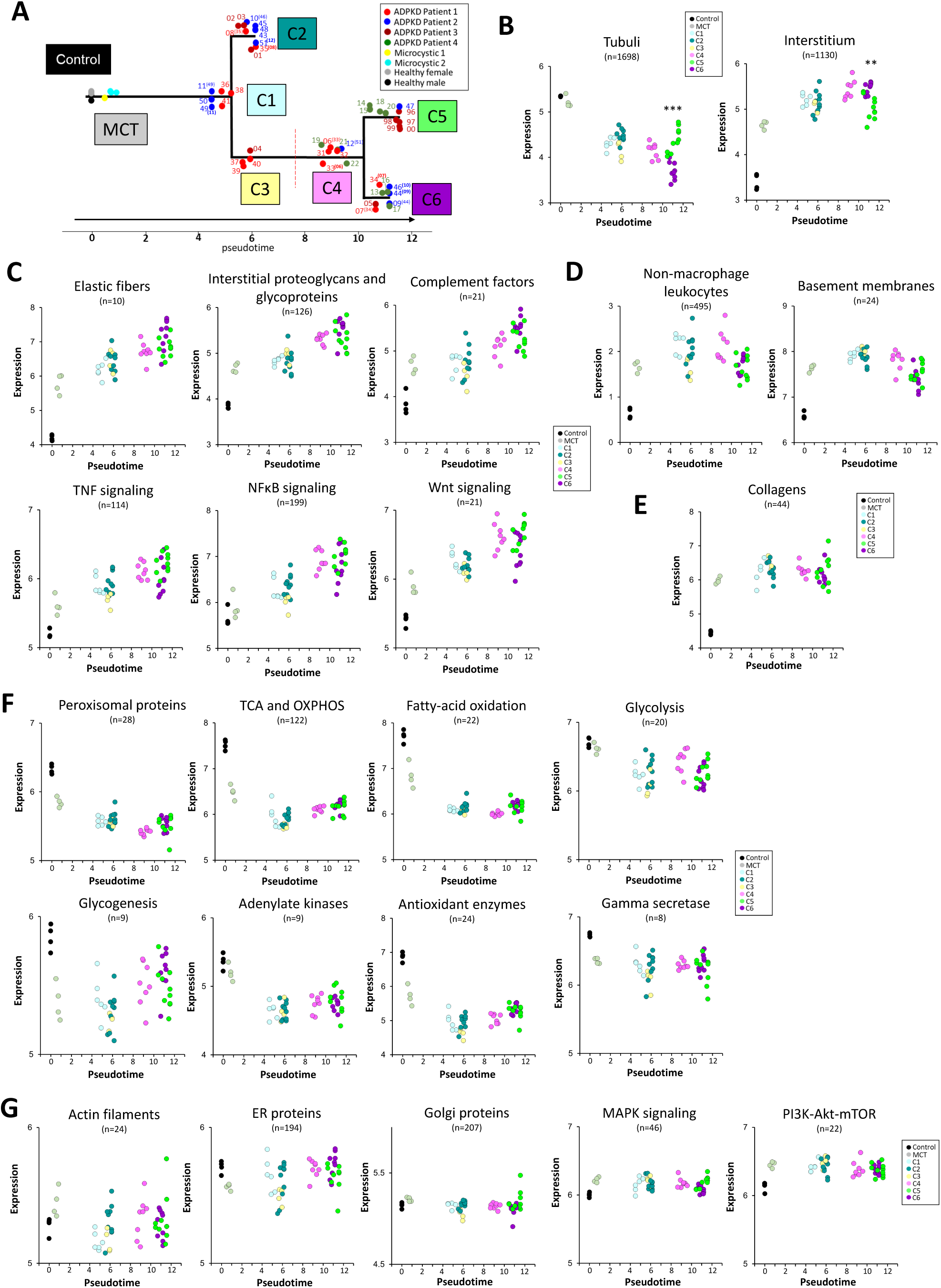
(previous page): Main trajectory in cyst samples from ADPKD patients. A) Trajectory reconstruction analysis of the different samples. The red dashed line indicates the separation of the 2 larger clusters identified in Fig. 1H. All samples are assigned by a number, while the color corresponds to the specific patients. For adjacent samples from the same cyst, the number of the other sample is indicated in brackets. The colors corresponding to the 6 subclusters are indicated in the rectangles C1-C6. B) Expression of all common renal tubular (left) and interstitial (right) markers over the trajectory pseudotime. C) Expression of different categories that showed a gradual increase over the trajectory pseudotime, without deviation of C2 and between C5 and C6. D) Expression of different categories that first showed a gradual increase, followed by a decrease over the trajectory pseudotime, without deviation of C2 and between C5 and C6. E) Expression of collagens is similarly increased in all subclusters, without deviation of C2 and between C5 and C6. F) Expression of different categories that showed a decrease over the trajectory pseudotime, without deviation of C2 and between C5 and C6. G) Expression of different categories that do not change over the trajectory pseudotime. n = the number of transcripts per category. **p<0.01, ***p<0.001 (linear mixed-model analysis).

We first characterized the categories that were similar in C2 and C3, and in C5 and C6, and that therefore showed no deviations along the trajectory. A gradual increase in expression was observed for elastic fibers, interstitial proteoglycans and glycoproteins, complement factors and TNF-, NFκB- and Wnt-signaling pathways (Fig. 2C). Non-macrophage leukocytes and basement-membrane expression was increased in MCT and C1-C3, followed by a decrease in C5 and C6 (Fig. 2D). Collagen expression was similarly increased in all clusters (Fig. 2E). Oppositely, a common downregulation was observed for expression of peroxisomal markers, metabolic pathways (tricarboxylic-acid (TCA) cycle and oxidative phosphorylation (OXPHOS), fatty-acid oxidation, glycolysis and glycogenesis), adenylate kinases, antioxidant enzymes and the gamma-secretase complex (Fig. 2F). The expression of actin filaments, ER proteins, Golgi proteins, the MAPK-signaling pathway and members of the PI3K-Akt-mTOR complexes did not significantly change along the trajectory (Fig. 2G and Suppl. Table 5).

Next, we characterized the deviations of C2, and between C5 and C6. Leaf-marker analysis with ClueGo-pathway-overrepresentation analysis (Suppl. Table 6) revealed that alterations in epidermis and epithelium development, and cell junctions defined the trajectory from C1 to C2. C4-C5 was associated with changes in TNF-related signaling and leukocyte differentiation, while C4-C6 was marked by changes in extracellular-matrix organization and the complement system (Suppl. Fig. 3). Furthermore, C5 showed a significant decrease compared to C6 in the expression of growth factors, TIMPs, macrophages, fibroblasts, endothelial cells, profibrotic transcripts and microtubules, and an increased expression of mitochondrial proteins, enzymes catalyzing xenobiotic detoxification, intercellular junctions and developmental proteins (Fig. 3A). C2 showed a significantly higher expression in ADPKD-associated transcription factors, apoptosis, dual oxidases, epithelial keratins and class I UDP-glucuronosyltransferases (UGTs) (Fig. 3B). Moreover, histological collagen analysis of the cystic membranes by Sirius-Red staining revealed that the collagen fibers of C2 were significantly less thick compared to those in other subclusters (Fig. 3C).

**Fig. 3:**
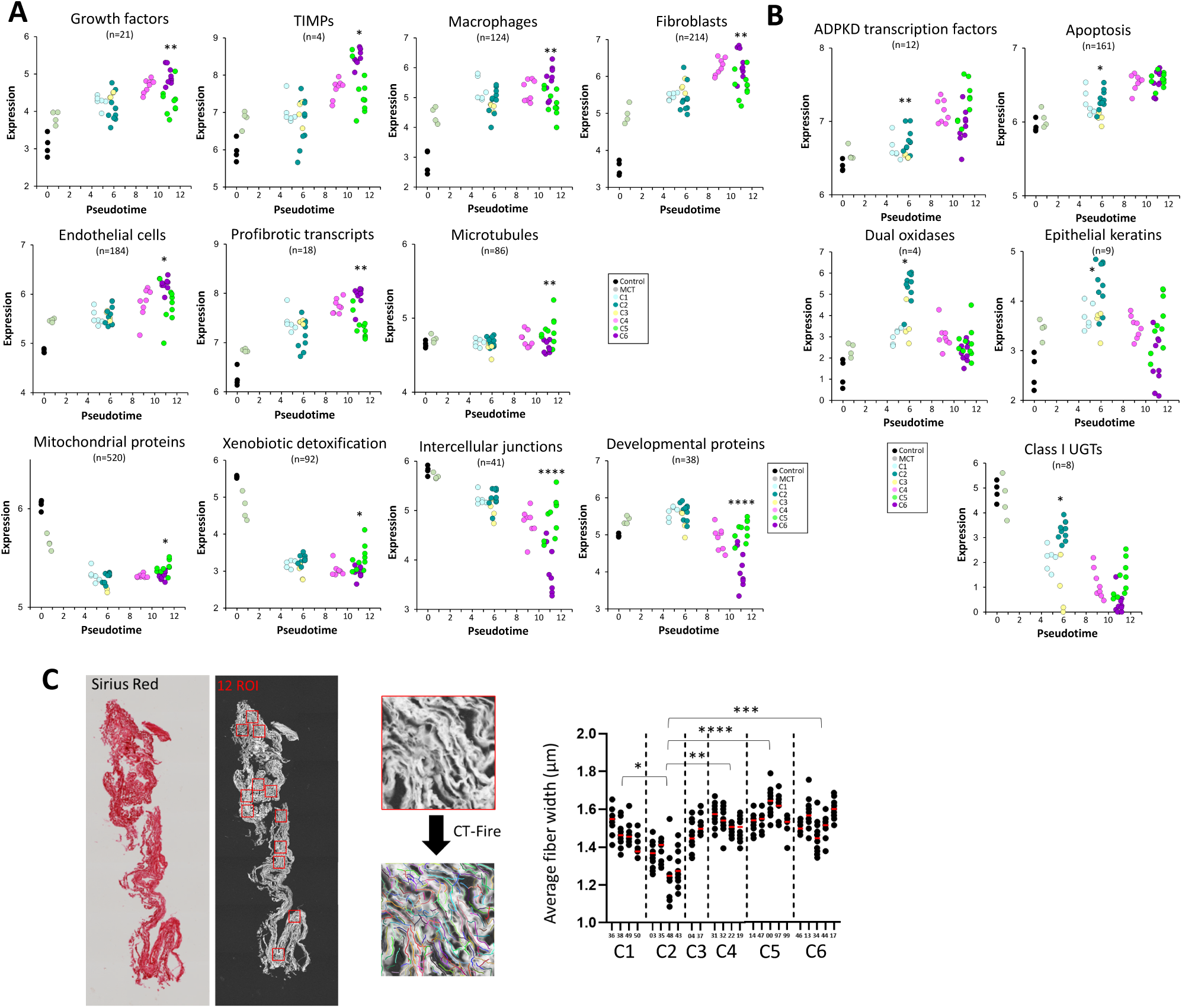
Deviating trajectory of C2 and between C5 and C6 in cyst samples from ADPKD patients. A) Expression of different categories over the trajectory pseudotime that showed a significantly different expression in C5 versus C6. B) Expression of different categories over the trajectory pseudotime that showed a significantly different expression in C2 versus C3. *p<0.05,**p<0.01,****p<0.0001 (linear mixed-model analysis). C) Histological analysis of collagen fibers. Left: example of a cystic membrane stained with Sirius Red, and the indication of 12 analyzed regions of interest (ROI). Middle: example quantification of the average fiber width in a ROI with the CT-Fire algorithm. Right: quantification of the average fiber width per region of interest of each sample of the 6 subclusters (C1-C6).*p<0.05,**p<0.01,***p<0.001,****p<0.0001 (One-Way ANOVA).

### Expression of tubular-segment markers

Specific markers for most tubular cell types showed a downregulation in the cyst samples, except for the thin ascending limb that showed an upregulation (Fig. 4A). All segment-specific transcripts showed a higher expression in C2 versus C3 and/or in C5 versus C6 (Fig. 4A). The downregulation was most pronounced for proximal tubular markers with already a significant decrease in the MCT, which did not occur for the other tubular-segment markers (Fig. 4A and Suppl. Table 5). In several cystic samples, the expression of non-proximal tubular markers was even comparable to the expression in healthy tissue samples. Interestingly, the same samples (cyst 99, 96, 97, 00 and 47 from C5; 03 and 48 from C2) showed a relatively high expression of markers of multiple tubular cell types (Fig. 4A).

**Fig. 4.**
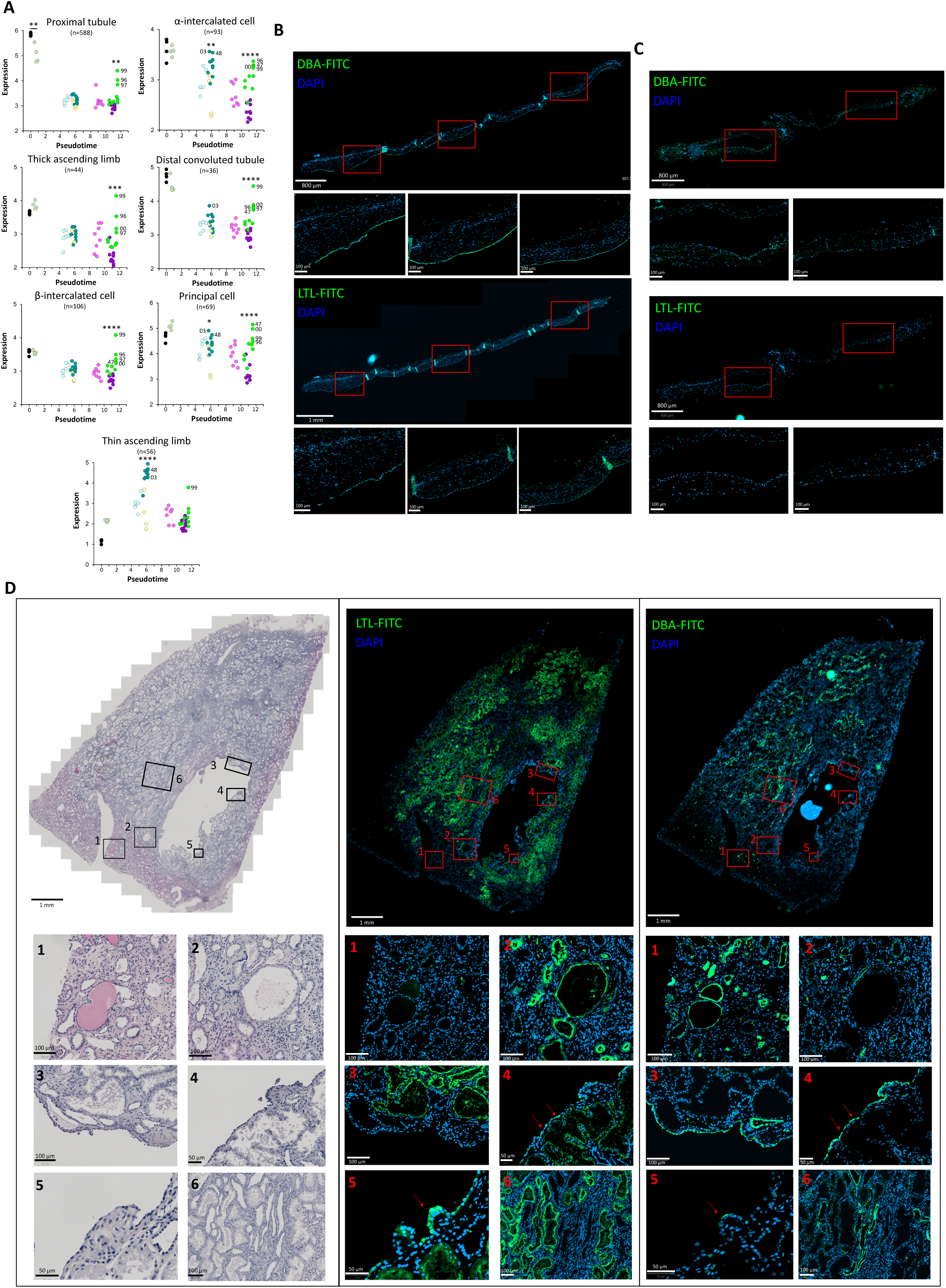
(previous page): Expression of tubular cell-type markers in macro- and microcysts from ADPKD patients. A) Expression of transcripts specific for different tubular cell types over the trajectory pseudotime. The colors of the data points correspond to the different subclusters of the trajectory analysis. Some samples are indicated by their designated number in all plots. B,C) Staining of the membrane of cyst 4G (B) and cyst 4D (C) with the collecting-duct marker DBA-FITC (upper) or the PTEC marker LTL-FITC (lower) and DAPI. For each staining the whole-tissue section and enlargements of the 3 insets with their respective scale bars are shown. D) Staining of a MCT with H&E (left), LTL-FITC and DAPI (middle) or DBA-FITC and DAPI (right). The whole-tissue section and enlargements of the 6 insets with their respective scale bars are shown. The red arrows indicate areas where both positive LTL and DBA staining occurred. *p<0.05,**p<0.01,***p<0.001,****p<0.0001 (linear mixed-model analysis).

This relative difference in downregulation between proximal and distal tubular markers was also evident from staining of the cyst samples with either Dolichos Biflorus (Horse Gram) Agglutinin-fluorescein (DBA-FITC) or Lotus Tetragonolobus (Asparagus Pea) Lectin-fluorescein (LTL-FITC), which targets collecting-duct cells or proximal tubular cells, respectively. In various cystic tissues, positive DBA staining could be observed in the cyst-lining cells, while LTL staining remained negative in all analyzed specimens (Fig. 4B-C). Nonetheless, we identified microcysts in MCT, which also contained normal renal parenchyma, glomeruli and tubules (Fig. 4D), that stained positive for DBA (Fig. 4D, inset 1) as well as for LTL (Fig. 4D, inset 2). In this tissue, a small macrocyst (with a diameter in the millimeter range), contained cyst-lining epithelium positive for DBA (inset 3-5) and LTL (inset 4-5), indicating that cysts could carry markers of different tubular cell types. Normal tubules never displayed double DBA- and LTL-positive staining (inset 6).

### Modulation of purine metabolism attenuates in vitro cyst development

Since cysts were heterogeneous, we focused on changes that were comparable in all ADPKD samples including MCT to identify potential targets to modulate cystogenesis in all cysts. Metabolic alterations were among these homogenous alterations (Fig. 2D). We therefore performed tracer metabolomics with ^13^C-labeled D-glucose in human kidney cell lines derived from the urine of young healthy individuals or early-stage ADPKD patients (proximal tubular epithelial cells (PTECs) or collecting-duct epithelial cells (CDECs); Fig. 5A) (Decuypere et al., 2021; Janssens et al., 2021), and in tissue-derived cell lines from healthy kidney tissue or cystic membranes (late-stage ADPKD). As previously reported for the urine-derived PTECs (Decuypere et al., 2021), the CDECs from young ADPKD patients have lower PC1 levels compared to CDECs from healthy controls (Fig. 5B). In line with the transcriptomic downregulation of TCA enzymes (Fig. 2D), the incorporation of D-glucose into the TCA-cycle metabolites was significantly reduced in urine-derived early-stage PTECs and in tissue-derived late-stage cystic cells from ADPKD patients, compared to cells from healthy individuals. However, labeling of TCA metabolites was similar in early-stage CDECs from ADPKD patients versus healthy individuals (Fig. 5C), suggesting that changes in TCA-metabolite labeling was segment dependent. We therefore searched for metabolomic changes in ADPKD versus healthy cells that occurred in all tubular cell types. We observed significantly enhanced m5 labeling specifically in the adenosine derivates AMP and cAMP in all human ADPKD tubular cell types (early-stage PTECs and CDECs, late-stage cystic cells), as well as in ADPKD murine polycystin-1 knockout (PC1KO) versus wild-type (WT) cell models, namely mouse inner medullary collecting-duct cells (mIMCDs) and mouse embryonic fibroblasts (MEFs) (Fig. 5D and Suppl. Fig. 4). The m5 labeling points to the labeling of the ribose in the nucleotide and suggests alterations in the pentose phosphate pathway (PPP) or purine metabolism (Verdegem *et al,* 2017). A change in PPP was unlikely, since the pyrimidine nucleotides CMP, CDP, CTP, UMP, UDP, UTP did not show enhanced m5 labeling (Fig. 5D and Suppl. Fig. 4). Additionally, changes in the PPP were not observed in the RNA-Seq of the human cyst samples, labeling of PPP metabolites was not enhanced in human and mouse ADPKD cell lines, and blocking the PPP with G6PDi-1 did not affect in vitro cyst formation in human cyst cell lines (Suppl. Fig. 5).

**Fig. 5:**
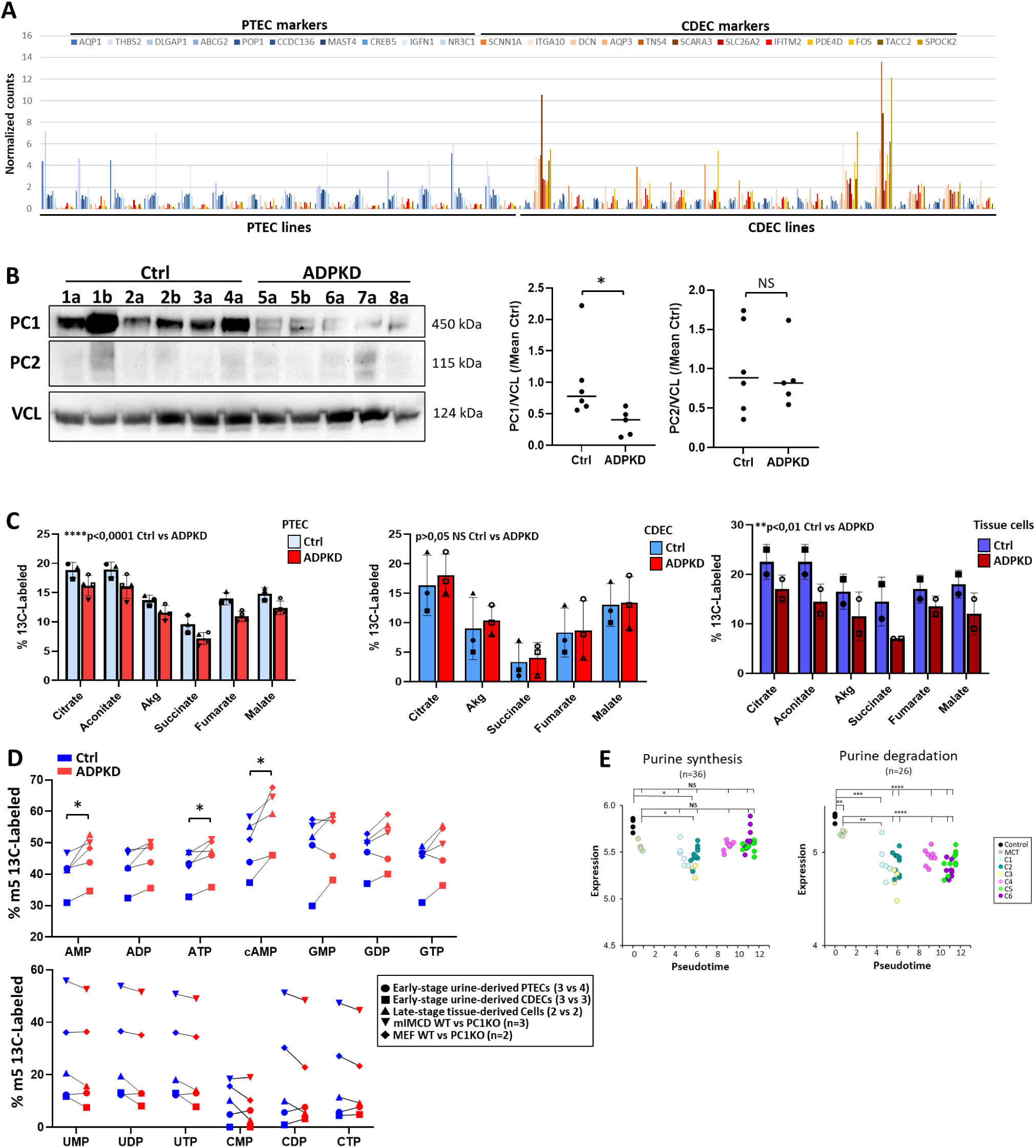
Altered purine metabolism in ADPKD cell lines and tissues. A) RNA-Seq-based normalized expression of selected PTEC (blue) or CDEC (orange) markers in 14 PTEC and 14 CDEC cell lines. Each group contained 7 individuals with 2 monoclonal cell lines each. B) Western-blot analysis of Polycystin-1 (PC1), -2 (PC2) and Vinculin (VCL) in urine-derived CDEC from young healthy individuals (Ctrl) and ADPKD patients. For each individual (indicated by the number 1-8), one (a) or two (a, b) monoclonal cell lines were analyzed. Left: immunoblot; right: quantification of PC1 or PC2 over VCL levels. * p < 0.05, NS not significant (Mann-Whitney test). C) Fractional contribution of ^13^C derived from ^13^C-D-glucose to the TCA metabolites in urine-derived early-stage PTECs (left), urine-derived early-stage CDECs (middle) and tissue-derived late-stage cells (right) from healthy individuals (blue) or ADPKD patients (red). **p<0.01, ****p<0.0001 (One-Way ANOVA). D) Fractional m5 labeling in adenosine nucleotides (AMP, ADP, ATP and cAMP), guanosine nucleotides (GMP, GDP, GTP), uridine nucleotides (UMP, UDP, UTP) and cytidine nucleotides (CMP, CDP, CTP) in control (blue) and ADPKD (red) cell models. The symbols represent the average values of urine-derived early-stage PTECs (3 controls and 4 ADPKD cells; circles), urine-derived early-stage CDECs (3 controls and 3 ADPKD cells; squares), tissue-derived late-stage cells (2 controls and 2 ADPKD cells; triangles), mIMCDs (WT versus PC1KO; n = 3; reversed triangles) and MEFs (WT versus PC1KO; n = 2; diamonds). *p<0.05, **p<0.01 (multiple paired t-tests). E) Expression of transcripts for purine synthesis and degradation over the trajectory pseudotime in cyst samples from ADPKD patients. The colors of the data points correspond to the different subclusters of the trajectory analysis. NS, not significant, *p<0.05,**p<0.01,***p<0.001,****p<0.0001 (linear mixed-model analysis).

Purine-degradation transcripts were significantly downregulated in all cyst subclusters and already occurred in MCT (Fig. 5E). Purine biosynthesis on the other hand was not significantly affected (Fig. 5E). To analyze whether purine metabolism played a role in cyst development, we used the purine-synthesis inhibitor azathioprine in an in vitro cyst assay, using 3 independent human cyst cell lines derived from cystic tissue of 3 nephrectomized patients. Azathioprine attenuated FSK/IBMX-induced cyst formation, and became effective at concentrations around 1 - 3 µM in all 3 cell lines (Fig. 6A and Suppl. Fig. 6A). In contrast to the effect of rapamycin, azathioprine did not significantly affect 7-day cyst-cell proliferation (Fig. 6B and Suppl. Fig. 6B), nor did it significantly alter the migration and proliferation of the cells around the Matrigel dome after 20 days in the in vitro cyst assay (Fig. 6C and Suppl. Fig. 6C). This suggest that purine-metabolism inhibition attenuated cyst growth, but not via proliferation or migration.

**Fig. 6:**
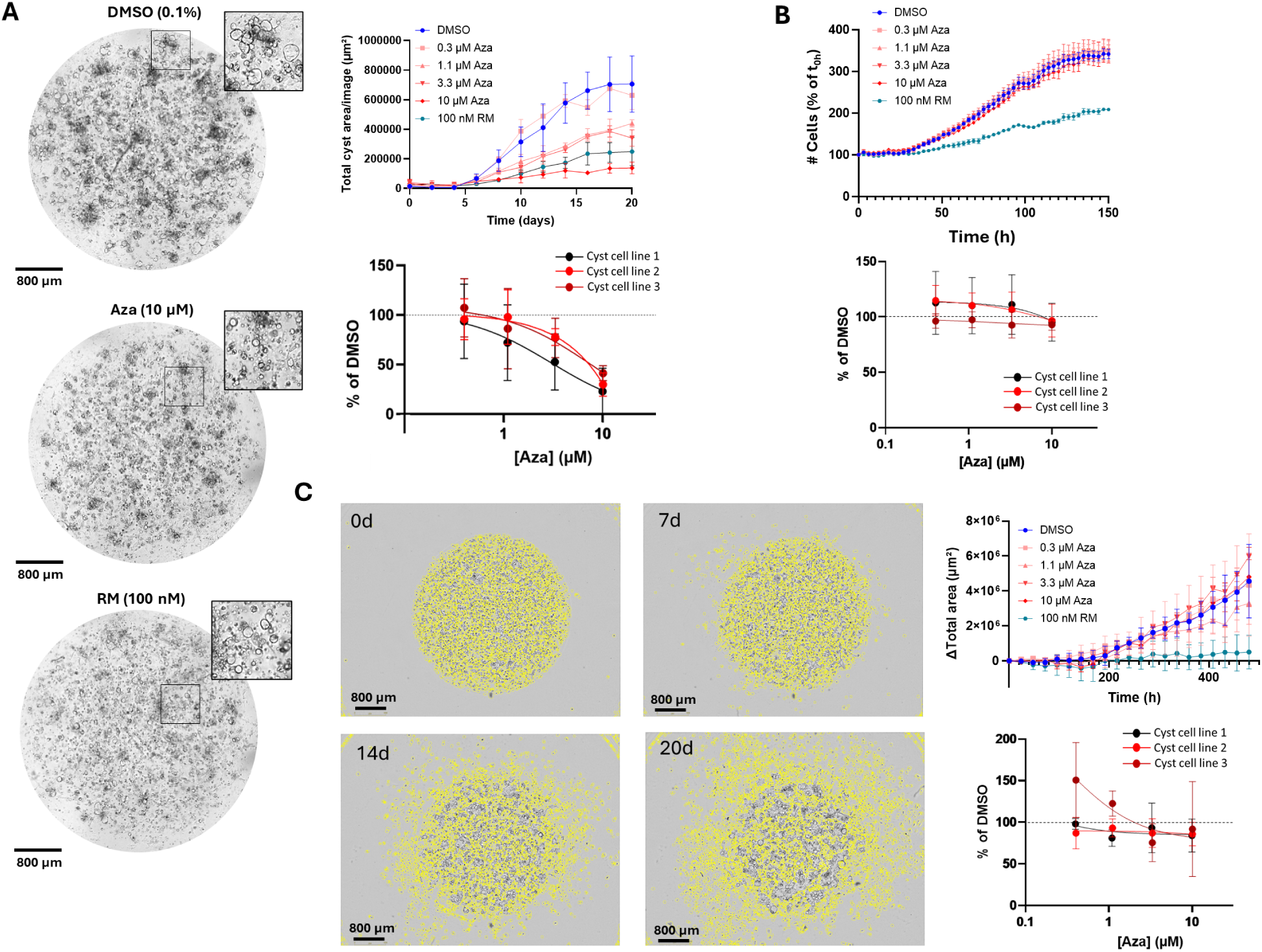
Azathioprine attenuates in vitro cyst formation. A) In vitro cyst assay of human cyst-tissue-derived cells in 50% Matrigel domes treated with DMSO, azathioprine (Aza) or rapamycin (RM). Left: representative images after 20 days of cyst growth; Upper right: quantification of total cyst area per image over time with different concentrations of azathioprine or 100 nM rapamycin in a representative cell line; Lower right: dose response for 3 independent cell lines (n = 3 per cell line) for azathioprine. B) Proliferation assay of human cyst-tissue-derived cells treated with various concentrations of azathioprine or 100 nM rapamycin. Upper: representative experiment; Lower: dose response for 3 independent cell lines derived from human cystic tissue (n = 4 per cell line). C) Outgrowth out of the Matrigel dome of human cyst-tissue-derived cells treated with various concentrations of azathioprine or 100 nM rapamycin. Left: representative images of the quantification of the outgrowth over time; Upper right: quantification of the outgrowth in a representative cell line in the presence of various concentrations of azathioprine (Aza) or of rapamycin (RM; 100 nM); Lower right: dose-response curves of 3 independent cell lines (n = 3 per cell line) for azathioprine.

## DISCUSSION

We performed RNA-Seq-based transcriptomics using samples from 40 individual kidney cysts from 4 ADPKD patients with *PKD1* mutations. Over 2500 genes were differentially expressed in the cystic compared to healthy tissue. However, the significance of these alterations is difficult to interpret since cystic tissue has a different cellular constitution than healthy kidney tissue. We therefore focused on the transcriptomic heterogeneity amongst the cyst samples, which was independent of patient, size and location of the cysts. In order to describe the observed heterogeneity, principal-component, cluster and trajectory analyses were performed. Cyst samples were grouped into 2 clusters and 6 subclusters and were heterogeneous within a single patient. In addition, adjacent membrane pieces of the same cyst did not always group together, suggesting that the membrane surrounding a single cyst was not per se uniform, and was composed of regions with different expression patterns.

Overall, the heterogeneity correlated with a decrease in tubular markers and an increase in interstitial markers (Fig. 2B). The observed heterogeneity cannot simply be explained by an increase in the thickness of the cyst walls, or of a higher proportion of interstitium versus cyst-lining cells. Some (Fig. 2C) but not all (Fig. 2D-E) aspects of the interstitium showed a gradual increase along the trajectory, while other interstitium-related categories significantly deviated between subclusters (Fig. 3A). In addition, differences in thickness of collagen fibers between the subclusters (Fig. 3C) pointed to structural remodeling of the extracellular matrix. Finally, cluster 2 (composed of the subclusters C4, C5 and C6) are marked by the highest expression of interstitium, and they are also more associated with signatures of kidney damage, fibrosis and cell death (Fig. 1I). All this suggests that the observed heterogeneity is partially explained by structural alterations in the interstitium promoting fibrosis and kidney-tissue injury. Indeed, the interstitial matrix is remodeled in ADPKD and fibrosis is a significant manifestation in ADPKD associated with the progression to kidney failure (Fragiadaki et al., 2020; Norman, 2011; Zhang et al., 2020).

Based on the above, we hypothesize that the pseudotime in the trajectory analysis represents an evolution in disease progression, with a gradual increase in fibrosis and inflammation, and a concomitant gradual decrease of tubular markers, reflecting dedifferentiation. However, two deviations were observed along the trajectory: one of C2 and a deviation between C5 and C6. These subclusters (mostly C5) deviated by less dedifferentiation (i.e. higher tubular markers) and reduced expression of specific interstitial components, despite following the general trend in other ADPKD-related gradual changes (e.g. signaling pathways, metabolic changes; Fig. 2). Interestingly, C2 has some very specific signatures (e.g. enhanced epithelial keratins, dual oxidases and class-I UGTs; Fig. 3B). It would be interesting to better understand the significance of these signatures and how they affect cyst formation or disease progression.

With the respect to the dedifferentiation, the expression pattern of tubular markers was different depending on the segment investigated. Most notably, the decrease of proximal tubular markers was more pronounced and already present in MCT, compared to the distal markers. In some cystic tissues, distal-marker expression was even comparable to healthy tissues. This was confirmed by a positive DBA staining of cyst-lining cells in certain cystic tissues, while none stained positively for LTL. However, positive LTL staining was observed in some microcysts, suggesting that cysts indeed originated from proximal tubular cells, but their dedifferentiation rate was more pronounced compared to other tubular segments.

Interestingly, the same cystic samples showed a relatively high expression of markers of various tubular segments (Fig. 4A). This is in line with a positive staining for both DBA and LTL in a small macrocyst (Fig. 4D). Such a simultaneous expression of multiple tubular segment markers in a cyst has also been previously described (Li et al., 2021). This heterogeneous composition can also explain the different transcriptomic profiles of two adjacent cystic membranes in the same cyst (see above). Although this is possibly due to fusion of individual smaller cysts (Li et al., 2021), the remarkable upregulation of thin-ascending-limb markers in all samples (Fig. 4A) may also indicate that during dedifferentiation cysts start to display aberrant expression patterns.

To evaluate potential targets to modulate cyst formation in ADPKD, we focused on common changes occurring in all cysts and all subclusters in a similar rate and without any deviations along the trajectory. Furthermore, significant changes already observed in MCT are more likely primary events early in cyst development. These general homogenous changes include an upregulation of leukocyte transcripts and collagens, as well as a downregulation of metabolic processes. The latter was therefore further explored experimentally via tracer metabolomics in human cell lines derived from urine of pediatric early-stage ADPKD patients or young healthy individuals, and from cystic tissue of late-stage ADPKD kidneys or from healthy kidney tissue. We confirmed a decrease in the TCA-cycle-metabolite labeling in early-stage PTECs and late-stage cystic cells, but not in early-stage CDECs. We hypothesize that this difference between PTECs and CDECs also reflects the different rates of dedifferentiation between the cell types.

As the labeling of TCA metabolites was dependent on the cell type, we searched for homogenous alterations in all cell lines that corresponded with homogenous changes in the transcriptomic analysis of the cyst tissues. In this manner, we identified an enhanced m5 labeling (reflecting labeling of ribose) in adenosine-derived nucleotides, and most prominently AMP and its derivate cAMP. As this enhanced labeling was not observed for pyrimidine nucleotides and less prominent in guanosine nucleotides, it suggests a change specific for adenosine nucleotides. In the transcriptomic analysis, this corresponded with a similar significant downregulation of purine degradation transcripts in all cysts and MCT, while no significant changes in purine synthesis were observed (Fig. 5E). Furthermore, the purine breakdown products xanthine and hypoxanthine were less abundant in kidney tissues of the PKD1(RC/RC) mouse model (Hopp et al., 2022). Also urinary metabolomics in conditional PKD1 knockout mice confirmed altered purine-metabolism activity (Menezes et al., 2012), confirming earlier findings in other polycystic kidney models (cystic jck mice and PCK rats) (Taylor et al., 2010). Interestingly, azathioprine, a general purine-synthesis inhibitor, was able to inhibit cyst growth at clinically relevant concentrations (Tapner et al., 2004). In contrast to rapamycin, it did not inhibit proliferation, suggesting another, still elusive, mechanism of action. Based on these data, we hypothesize that ADPKD cells have an altered purine metabolism, already present in the early stages of the disease, which can be targeted to reduce cyst formation.

Purine analogues such as azathioprine (Imuran®) are a common clinical therapy to treat autoimmune disorders (e.g. Crohn’s disease and rheumatoid arthritis), to prevent rejection after transplantation (Díaz-Villamarín et al., 2023), and with potential towards cancer therapy (Yin et al., 2018). Although it is generally used as an immunosuppressant, it only affects rapidly proliferating (immune or cancer) cells that are dependent on de novo purine synthesis, leaving differentiated epithelial cells, who rely mostly on purine salvage, mostly undisturbed (Maltzman & Koretzky, 2003). However, as our data suggest, the effect of purine-synthesis inhibition by azathioprine on cyst cells could be independent of its effect on proliferation. As components of various molecules, purines have a central role in cell proliferation (DNA), metabolism (e.g. ATP, NAD(P)H, coenzyme A) and signaling (GDP, GTP, cAMP). The effect of azathioprine on all these aspects and how it affects the cellular phenotype and cystogenesis in ADPKD cells should be elucidated in future work.

In conclusion, through transcriptomic analysis of individual cysts, we tracked down the disease progression of cysts and their environment, and described which molecular signatures deviated from this pattern. By focusing on common alterations that already occurred at an early-stage, we identified altered purine metabolism as an important hallmark in ADPKD, which can be targeted to reduce cyst formation. Considering the current clinical application of purine-metabolism modulation for other diseases, its potential in repurposing for ADPKD treatment needs to be considered.

## METHODS

### Sex as a biological variable

All groups of the collected human samples and cell lines were contained male and female participants. Similar findings are reported for both sexes.

### Tissue collection

Sections of cystic membrane were dissected from polycystic kidneys from 4 nephrectomized genotyped patients (2 male and 2 female) with a confirmed *PKD1* mutation (Suppl. Table 1). From each patient, 9-11 cysts were analyzed resulting in a total of 40 cysts dissected (cyst 1A-K; cyst 2A-I; cyst 3A-J; cyst 4A-J; Suppl. Table 1). From 7 cysts, a second section was also analyzed, resulting in 47 samples in total. Whenever possible, additional sections were taken for histological analysis. Cyst size, region (cortical/medullary) and cyst-fluid color were notated (Suppl. Table 1). Four MCT samples, defined as macroscopically normal, were isolated from 2 other ADPKD patients with a confirmed *PKD1* mutation. As control, kidney parenchymal samples were taken during living-donation kidney transplantation from healthy donors (Suppl. Table 2). For RNA extraction, all tissues were snap-frozen in liquid nitrogen and stored at -80°C.

### RNA extraction and RNA-Seq

Following RNA extraction using the RNeasy Plus Mini kit (Qiagen) according to the manufacturer’s protocol, RNA concentration and quality were determined using DropSense96 (Perkin Elemer) and Bioanalyzer (Agilent). For RNA-Seq library preparation, the NEBNext® Ultra II Directional RNA Library Prep Kit for Illumina (Bioké) was used according to the manufacturer’s protocol. After quality assessment of the obtained libraries, an equimolar pool was prepared. A qPCR was then carried out using the Kapa SYBR FAST Universal qPCR kit for Illumina (Roche). Depending on the number of reads/samples required, the libraries were sequenced on a HiSeq4000 (Illumina). Quality control of raw reads was performed with FastQC v0.11.7 (http://www.bioinformatics.babraham.ac.uk/projects/fastqc). Adapters were filtered with ea-utils fastq-mcf v1.05 (https://github.com/ExpressionAnalysis/ea-utils). Splice-aware alignment was performed with HISAT2 (Kim et al., 2019) against the reference genome hg38 using the default parameters. Reads mapping to multiple loci in the reference genome were discarded. Resulting binary alignment map files were handled with Samtools v1.5 (Li et al., 2009). Quantification of reads per gene was performed with HT-seq Count v0.10.0, Python v2.7.14 (Anders et al., 2015). The data were normalized by using the variance stabilization transformation.

### Differential expression analysis

This analysis was performed using the data matrix of count data after filtering genes for a minimum of 10 counts in total including all samples (sum > 10) and selecting only protein-coding genes using the approved gene symbol from HGNC (https://www.genenames.org/) before normalization. Normalization was performed using counts per million (CPM) of gene counts controlling the normalization influence of highly expressed genes over the samples by subsetting to the 95% of cumulative expression (Weinreb et al., 2018). Normalized values were further transformed as pseudocounts using log_2_(CPM+1). Differential expression statistics were calculated based on a ranked *t*-test of each cluster (see cluster analysis below) against the healthy control cluster as reference, with the *p*-value adjusted by multiple testing (Bonferroni).

### Cluster analysis

Sample clustering was performed using the Leiden algorithm (Traag et al., 2019) with resolution 0.8 and visualization performed on dimensionality reduction by UMAP (Uniform Manifold Approximation and Projection), using nearest neighbor graph (McInnes L, 2018) with the number of neighbors of 10, overall identifying 3 major clusters (healthy, MCT and cysts) and 2 subclusters in the “cysts” cluster.

### Trajectory analysis

The reconstruction of cyst evolutionary trajectories was based on their similarity of transcriptomic information and was performed using STREAM following the standard published workflow with default parameters (Chen et al., 2019; Sprooten et al., 2024).

### Pathway analysis

For overrepresentation analysis the Cytoscape plug-in ClueGo was used (Bindea et al., 2009), while (comparative) expression analyses were performed in Ingenuity Pathway Analysis (Qiagen). Findings were only included when Benjamini-Hochberg-adjusted *p*-values were < 0.01 and absolute Z-scores > 2.

### Gene sets

To test for signatures, we created a panel of gene sets for each category relevant to ADPKD (Suppl. Table 3). For this purpose, transcripts for macrophages and non-macrophage leukocytes, fibroblasts, endothelial cells, epithelial cells and epithelial keratins from the different tubular segments were identified using the Kidney Interactive Transcriptomics database (http://humphreyslab.com/SingleCell/search.php) (Muto et al., 2022; Wu et al., 2018) and the Human Protein Atlas (https://v22.proteinatlas.org) (Karlsson et al., 2021). The Human Protein Atlas was also used to select growth factors, transcripts for the TCA cycle, OXPHOS and fatty-acid oxidation, genes encoding transcripts targeted to specific subcellular compartments and genes involved in the detoxification of xenobiotics. Only transcripts for which the subcellular expression has been experimentally verified with an approved, enhanced or supported reliability score were included. In addition to the Human Protein Atlas, literature was consulted for transcripts for elastic fibers (Heinz, 2021), antioxidant enzymes (Black et al., 2011), gamma-secretase subunits (Zhou et al., 2006), adenylate kinases (Fujisawa, 2023), profibrotic transcripts (Colon et al., 2019; Formica & Peters, 2020; Norman, 2011), TIMPs (Fragiadaki et al., 2020), dual oxidases (Faria & Fortunato, 2020), class-I UGTs (Meech et al., 2019), Wnt signaling (Lal et al., 2008), basement membranes (Zaferani et al., 2014) and ADPKD transcription factors (Zhou & Torres, 2022). For interstitial proteoglycans and glycoproteins (Hynes & Naba, 2012), the following categories were included: major known extracellular matrix glycoproteins, nervous system-enriched extracellular matrix proteins, vascular extracellular matrix proteins, extracellular matrix proteins of bones, cartilage and teeth, and cellular communication network factors. Kidney developmental genes were selected after a literature search (Berry et al., 2011; Brzóska et al., 2016; Fanni et al., 2011; Gill & Rosenblum, 2006; Harris et al., 2022; Jenkins et al., 2005; Kohl et al., 2014; Kunimoto et al., 2017; Lipp et al., 2021; Lokmane et al., 2010; Marcotte et al., 2014; Sonnenberg et al., 1991; Srinivas et al., 1999; Takamiya et al., 2004; Torban & Sokol, 2021; van der Ven et al., 2018; Wang et al., 2018; Wanner et al., 2019). Apoptosis, TNF-signaling, NFκB-signaling, purine-biosynthesis and -degradation transcripts were based on hallmark gene sets from the Molecular Signatures Database.

### Histological analysis

After 24 h fixation in 4% paraformaldehyde, tissue samples were processed and embedded in paraffin blocks. Sections (4 µm) were stained with hematoxylin & eosin (H&E) or Sirius Red according to standard operating procedures. On light microscopy, the cystic tissue samples showed cysts with characteristics of the cortex and/or the medulla (Suppl. Fig. 1). Staining for DBA-FITC and LTL-FITC occurred according to the manufacturer’s protocol (ThermoFischer Scientific, L32474 and L32480). Images of stained sections were acquired using a Zeiss AxioScan 7 microscope slide scanner and processed using QuPath. For analysis of collagen fibers, 12 randomly chosen regions of interest of the same size (512 × 512 px² with 1 px² = 0.1721 µm²) of the Sirius Red-stained images were chosen and analyzed using the CT-Fire v2.0 beta fiber detection Matlab plug-in (Bredfeldt et al., 2014).

### Mouse cell lines

The generation and characterization of WT or PC1KO mIMCDs (a kind gift from Dr. S Somlo and Dr. Y Cai, Yale School of Medicine, CT, USA), were described before (Decuypere et al., 2021). WT or PC1KO MEFs were a kind gift from Dr. A Boletta (San Raffaele Scientific Institute, Milan, Italy) (Distefano et al., 2009).

### Human cell lines

Human cell lines include PTECs and CDECs generated from urine or tissue of healthy individuals or genotyped ADPKD patients with a heterozygous frameshift mutation in *PKD1*. Cystic cells were derived from cystic membranes of nephrectomized PKD1 ADPKD kidneys. Urine-derived cells were from young individuals and early-stage patients (age 5-29 y) and tissue-derived cells from older persons and late-stage patients (age 47-65 y).

Urine-derived PTECs were collected, subcloned and expanded as previously described (Decuypere et al., 2021). In short, a pellet from centrifuged urine was grown in supplemented DMEM:F-12. Urine-derived CDECs were collected in the same manner, but the cell pellet was resuspended in a 1:1 mix of Renal Epithelial Cell Growth Medium 2 (PromoCell C26030) supplemented with Growth Medium Supplement Mix (C39606), 5% fetal bovine serum (FBS) and 100 U Pen/Strep, and DMEM high glucose (Gibco 31966-047) supplemented with 1% non-essential amino acids (Gibco 11140-035), 5 ng/ml bFGF (Peprotech 100-18B), 5 ng/ml PDGF-AB (Peprotech 100-00AB), 5 ng/ml EGF (Peprotech AF-100-15), 10% FBS and 100 U Pen/Strep. Primary CDECs were selected by fluorescence-activated cell sorting with an antibody against the collecting-duct marker L1CAM (Abcam ab95694) (Debiec et al., 1998).

Healthy tissue-derived PTECs and CDECs were isolated from biopsies during a living donor kidney transplantation and cystic tissue-derived cells from cystic membranes of ADPKD kidneys. Healthy tissue was cut and treated with collagenase D (0.67 mg/ml) for 1.5 h, while cystic membranes were subjected to trypsin treatment for 30 min. The treated tissues were then filtered using 125 µm (PTECs, cystic cells) or 90 µm (CDECs) sieves. After centrifugation at 260 × g for 7 min, pellets were added to a Percoll gradient (density 1.07 g/ml and 1.04 g/ml) and centrifuged at 1620 × g for 25 min with low acceleration and without brake. The layer between the Percoll fractions was then collected, washed and incubated in PTEC or CDEC medium.

Primary cells were conditionally immortalized with a retroviral construct containing SV40_LT_tsA58. This temperature-sensitive SV40 large T (LT) antigen makes SV40 LT unstable at 37°C. Hence, conditionally immortalized cells were cultured at 33°C. For experiments, monoclonal cell lines were incubated for 10 days at 37°C, which initiated cell differentiation (Decuypere et al., 2021). All cell lines were frequently tested for mycoplasma contamination.

### Tracer metabolomics

DMEM:F12 without glucose, HEPES and glutamine (Biowest L0091) was supplemented with 15 mM HEPES, 2.5 mM glutamine and 17.5 mM D-glucose or ^13^C-labeled D-glucose (Cambridge Isotopes CLM-1396). After 10 days at 37°C, differentiated cells were incubated with medium containing D-glucose or ^13^C-D-glucose for 24 h. After washing with 0.9% ice-cold NaCl solution, cells were scraped in ice-cold cellular extraction buffer (80% methanol containing 2 µM d27-myristic acid), centrifuged at 20000 × g for 15 min at 4°C and the supernatant was analyzed with liquid chromatography–mass spectrometry. Mass spectrometry (MS) measurements were performed using a Dionex UltiMate 3000 LC System (Thermo Scientific Bremen, Germany) coupled via heated electrospray ionization to a Q Exactive Orbitrap to a Q Exactive Orbitrap mass spectrometer (Thermo Scientific). Ten μl sample was taken from an MS vial and injected onto a 15 cm C-18 column (Acquity UPLC -HSS T3 1. 8 μm; 2.1 x 150 mm, Waters). A step gradient was carried out using solvent A (10 mM TBA and 15 mM acetic acid in milliQ) and solvent B (100% methanol). The gradient started with 5% of solvent B and 95% solvent A and remained at 5% B until 2 min post-injection. A linear gradient to 37% B was carried out until 7 min and increased to 41% until 14 min. Between 14 and 26 minutes the gradient increased to 95% of B and remained at 95% B for 4 minutes. At 30 min, the gradient returned to 5% B. The chromatography was stopped at 40 min. The flow was kept constant at 0.25 mL/min and the column was kept at 40°C throughout the analysis. The HESI-source operated at negative polarity mode using a spray voltage of 4.8 kV, sheath gas at 40, auxiliary gas at 10, the latter heated to 260°C. The ion transfer capillary temperature was 300°C. The mass spectrometer operated in full scan (range [70.0000-1050.0000]) and AGC target was set at 3.0E^+006^ using a resolution of 140000. Data collection was performed using the Xcalibur software (Thermo Scientific). The data analyses were performed by integrating the peak areas (El-Maven - Elucidata). To the protein pellet, 200 mM NaOH was added and incubated at 95°C for 30 min. After centrifugation (5000 rpm, 4°C, 10 min), the supernatant was used to determine the protein concentration with the bicinchoninic acid assay (Pierce). This protein concentration was used for normalization of the total abundance of the metabolites.

*3D Cyst assay-* Cystic cells were seeded in a 10 µl dome of 50% Matrigel (Corning 356232) at a density of 10000 cells per dome, and the gel allowed to solidify for > 30 min. PTEC medium with azathioprine (0.37 - 10 µM), G6PDi-1 (10 µM, Cayman Chemical) or DMSO (0.1%) was added on top. After 4 days at 33°C, 1 µM forskolin (FSK) and 10 µM IBMX were added additionally. Medium and compounds were refreshed twice a week. Images were acquired using the Organoid module in the IncuCyte S3 (Sartorius). For image analysis, NIS-Elements (5.30, Nikon Instruments Europe B.V.) was used. To identify cysts in brightfield images, we used Segment.ai, an AI module that is a part of the NIS.ai suite, a commercially available (licensed) software. We trained this neural network using a set of manually annotated images. Additional morphology and size filtering steps were applied to only detect closed, circular objects. The neural network was then used for counting the cysts and quantifying their size.

### Cell proliferation measurements

Cystic cells were seeded at 5000 cells per well and incubated at 33°C in an IncuCyte S3 to acquire images every 2 hours. Images were afterwards analyzed using the Cell-by-Cell module in the associated IncuCyte software to determine the cell count.

### NADPH measurement

Following incubation with G6PDi-1 (10 µM), NAPDH levels were determined using the NADP/NADPH-Glo Assay kit (Promega) according to the manufacturer’s protocol.

### Statistical analysis

To analyze the differences between the different subclusters, a linear mixed-model analysis with a random effect of cyst was used on transformed data (inverse hyperbolic sine, which handles a right-skewed distribution and deals with the presence of zero-values) with Tukey adjustments for multiple testing. The residual variability was allowed to differ between the subclusters. Pearson’s correlation analysis was also performed on the transformed data with GraphPad Prism 10.2.2.

### Study approval

Informed consent was obtained from all participants and collection of human material was approved by the Ethical Committee of the University Hospitals Leuven (S51837, S53364 and S61154).

### Data availability

The supporting data values can be found in the supplementary material. The RNA-Seq data of the human kidney (cyst) tissues is found on NCBI’s Gene Expression Omnibus: GSE289843.

## Supporting information

Supporting Data Values

## ACKNOWLEDGEMENTS

We would like to thank Sara Kerselaers, Lotte Vanmeerbeek and Nele Van Ranst for technical assistance; the Genomics Core at the University Hospitals Leuven for the sequencing and initial analyses; Dr. Steve Somlo, Dr. Yiqiang Cai and Dr. Ke Dong (Yale University, New Haven, CT, USA) for the mIMCD cells; Dr. Alessandra Boletta and Dr. Laura Cassina (San Raffaele Scientific Institute, Milano, Italy) for the MEF cells.

ADG is supported by Research Foundation Flanders (FWO) (Fundamental Research Grant, G0B4620N; Excellence of Science/EOS grant, 30837538, for ‘DECODE’ consortium; Strategic Basic Research or SBO grant, S000523N), KU Leuven (C1 grant, C14/24/122; and C3 grants, C3/23/067, C3/21/037, C3/22/022), Come Up Against Cancer or Kom op Tegen Kanker (KOTK/2018/11509/1 and KOTK/2019/11955/1), VLIR-UOS (iBOF grant, iBOF/21/048, for ‘MIMICRY’ consortium), Olivia Hendrickx Research Foundation (OHRF), and European Union (EU) Mission Cancer grant for the GLIOMATCH consortium (Project no. 101136670). DM and RV are supported by the Research Foundation Flanders (FWO) under the grant G0C8920N DM and JPD under the grant G060623N. DM is also supported under the clinical senior research grant 1804123N. DM and BB are members of the European Reference Network for Rare Kidney Diseases (ERKNet)– Project ID No 739532.

## AUTHOR CONTRIBUTIONS

JPD, DjM and RV designed the research study; JPD conducted the experiments; DiM, JP and BB provided material; JPD and BG acquired data; JPD, DM, PK, LM, SF, NC and HDS analyzed the data; PK and TR provided reagents; JPD, DM, PK, LM, HDS, BG, ADG, DM and RV wrote the manuscript.

## THE PAPER EXPLAINED

PROBLEM: Autosomal dominant polycystic kidney disease (ADPKD) is the most common inherited kidney disease, characterized by the progressive growth of kidney cysts and kidney enlargement leading to kidney failure. As such, it is responsible for 10% of the European dialysis population, with currently no curative treatment available. Despite the identification of various molecular pathways that are altered in ADPKD cells, the exact sequence of events and their relative importance remain unclear.

RESULTS: We have analyzed 40 individual cysts from 4 ADPKD patients and found that cysts are heterogeneous, even when they are from the same kidney. This heterogeneity was dependent on the progression of the disease with clear correlations with the rate of fibrosis, inflammation and (de)differentiation. In this way, disease progression can be tracked down on the level of the individual cyst. Next to the heterogeneity, we focused on common alterations in all cysts. By combining the analyses of the individual cysts with experiments in human and mouse ADPKD cell models, we identified altered purine metabolism as consistently altered in all ADPKD cysts and cell types. Purine metabolism modulation with azathioprine suppressed in vitro cyst formation, highlighting the importance of this phenomenon in ADPKD cystogenesis.

IMPACT: Our data describes the variability among individual cysts, dependent on ADPKD progression. This aids in determining the relative importance of molecular alterations in ADPKD. As a proof-of concept, we identified purine metabolism as a common altered pathway in all ADPKD cysts and cell types, which was previously undescribed in this context. Finally, we identified a drug acting on purine metabolism with potential for repurposing for ADPKD.

