## Supplementary figures and images for "The molecular landscape of polycystic kidneys is marked by common alterations in purine metabolism"

### Supplemental Figure 1

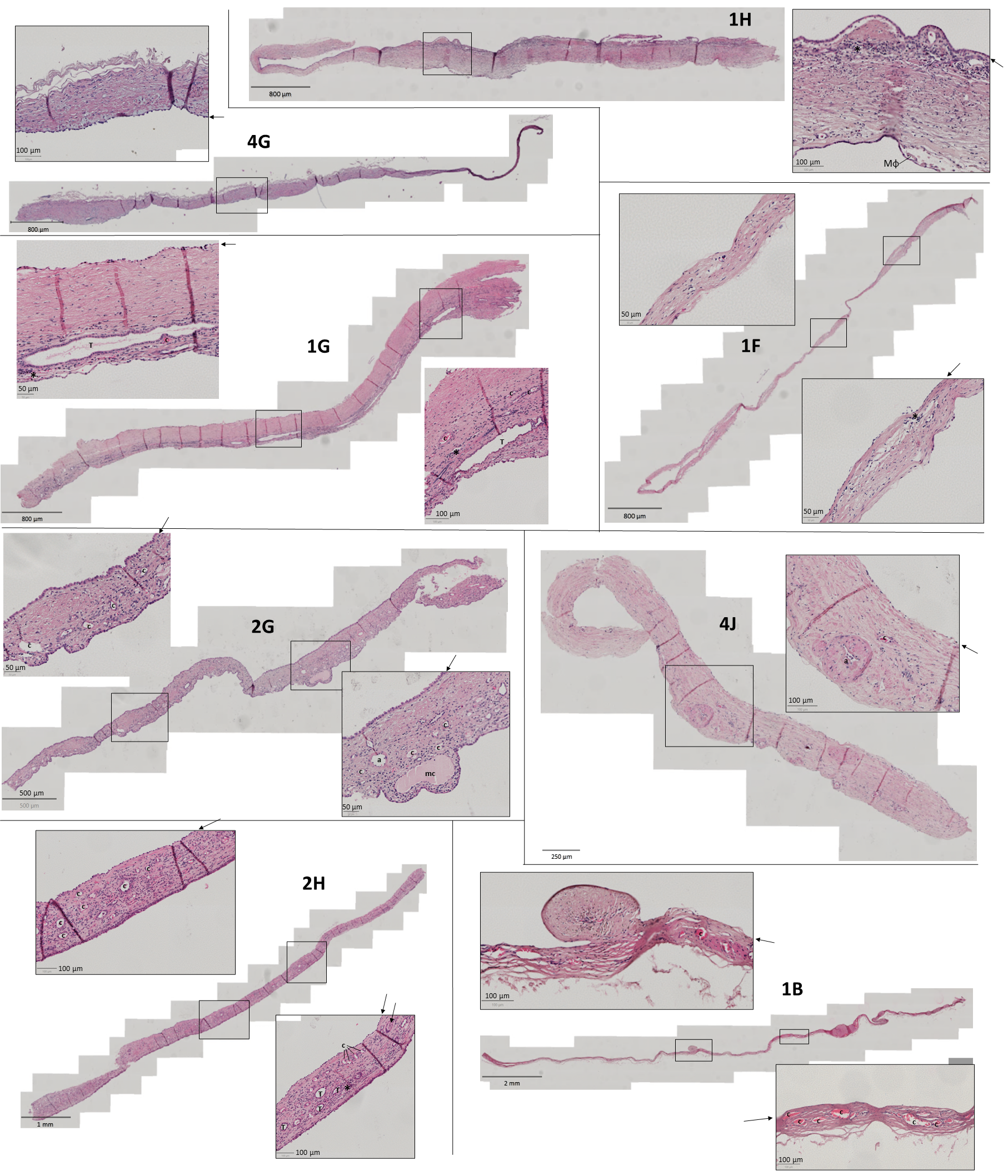

### Supplemental Figure 2

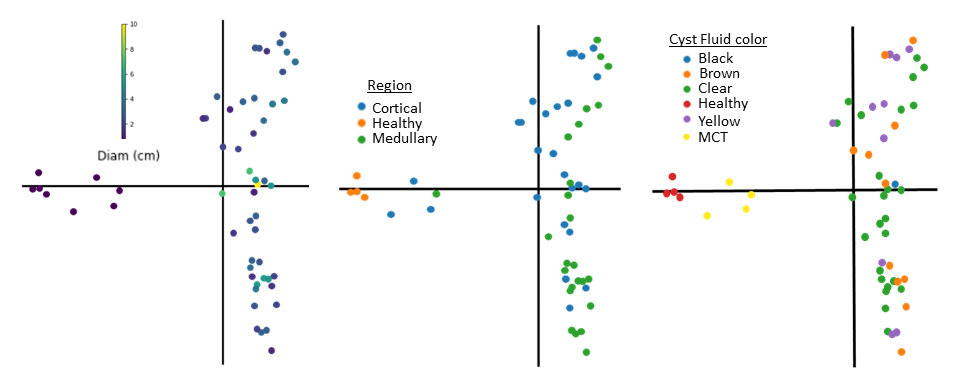

### Supplemental Figure 3

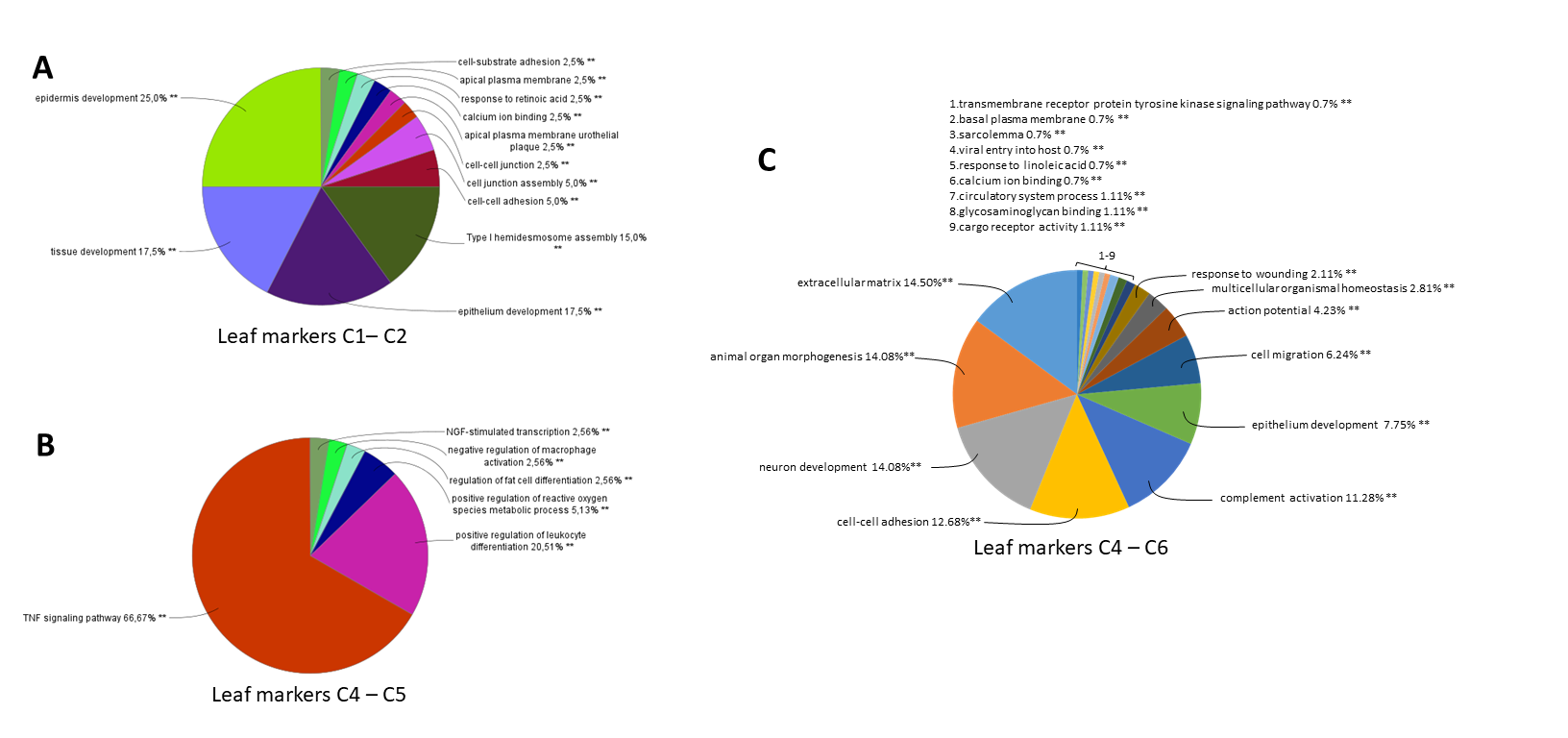

### Supplemental Figure 4

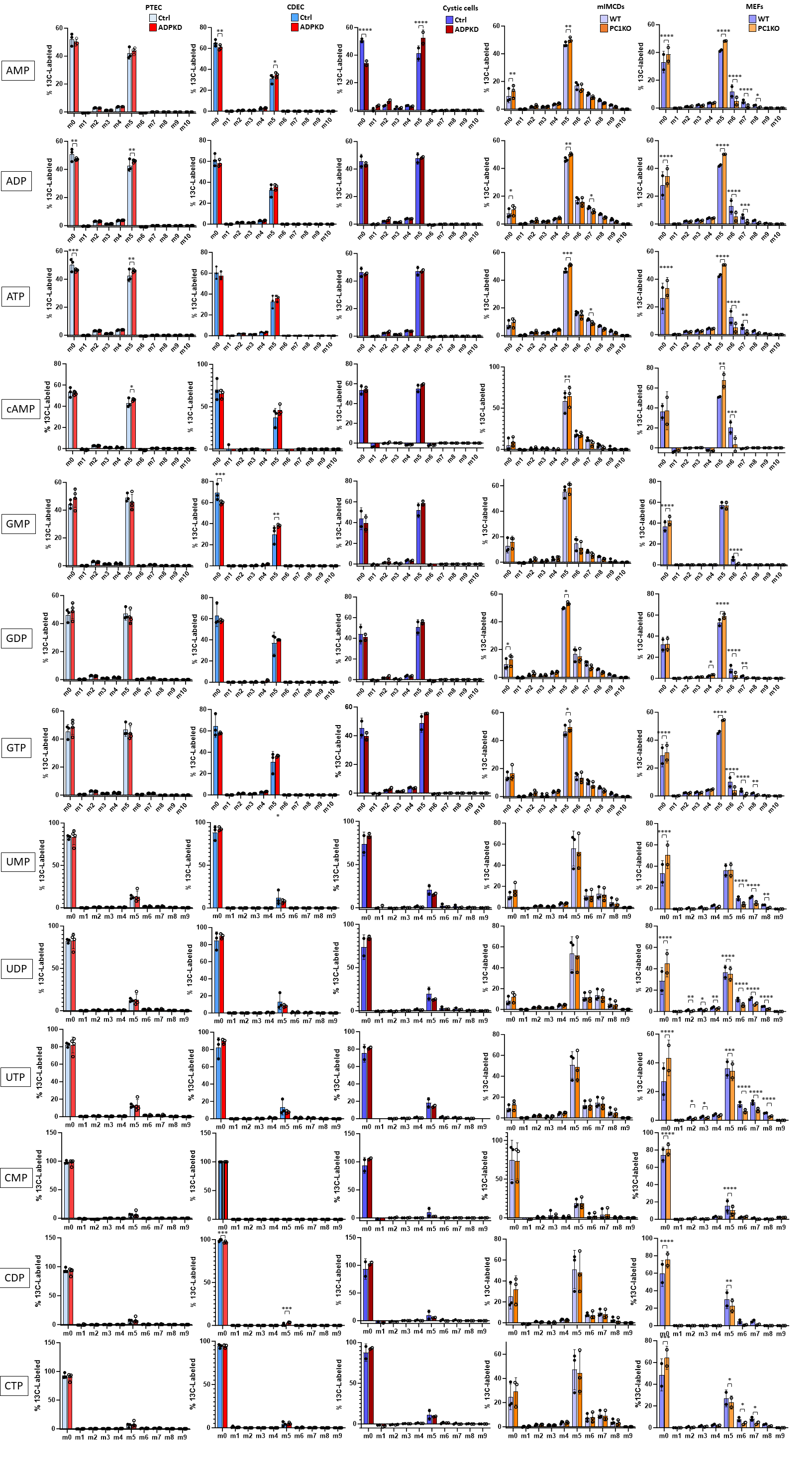

### Supplemental Figure 5

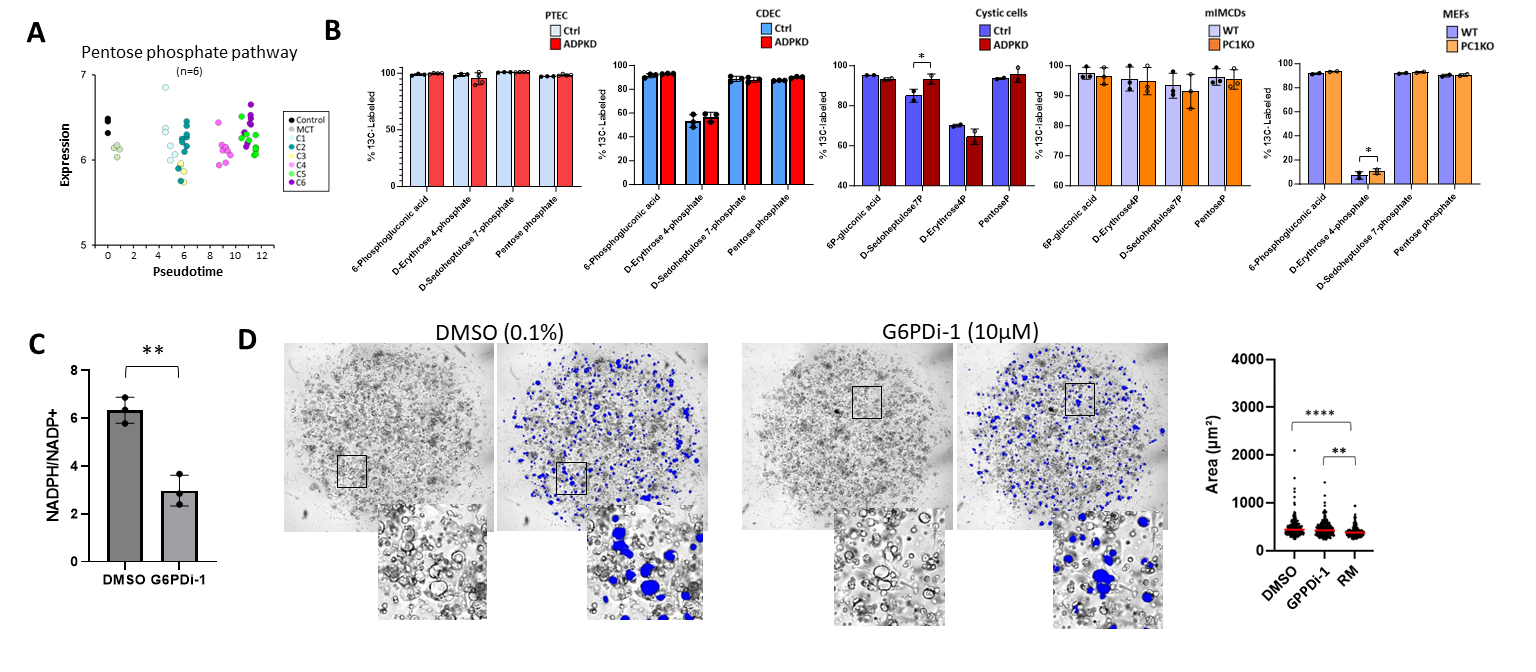

### Supplemental Figure 6

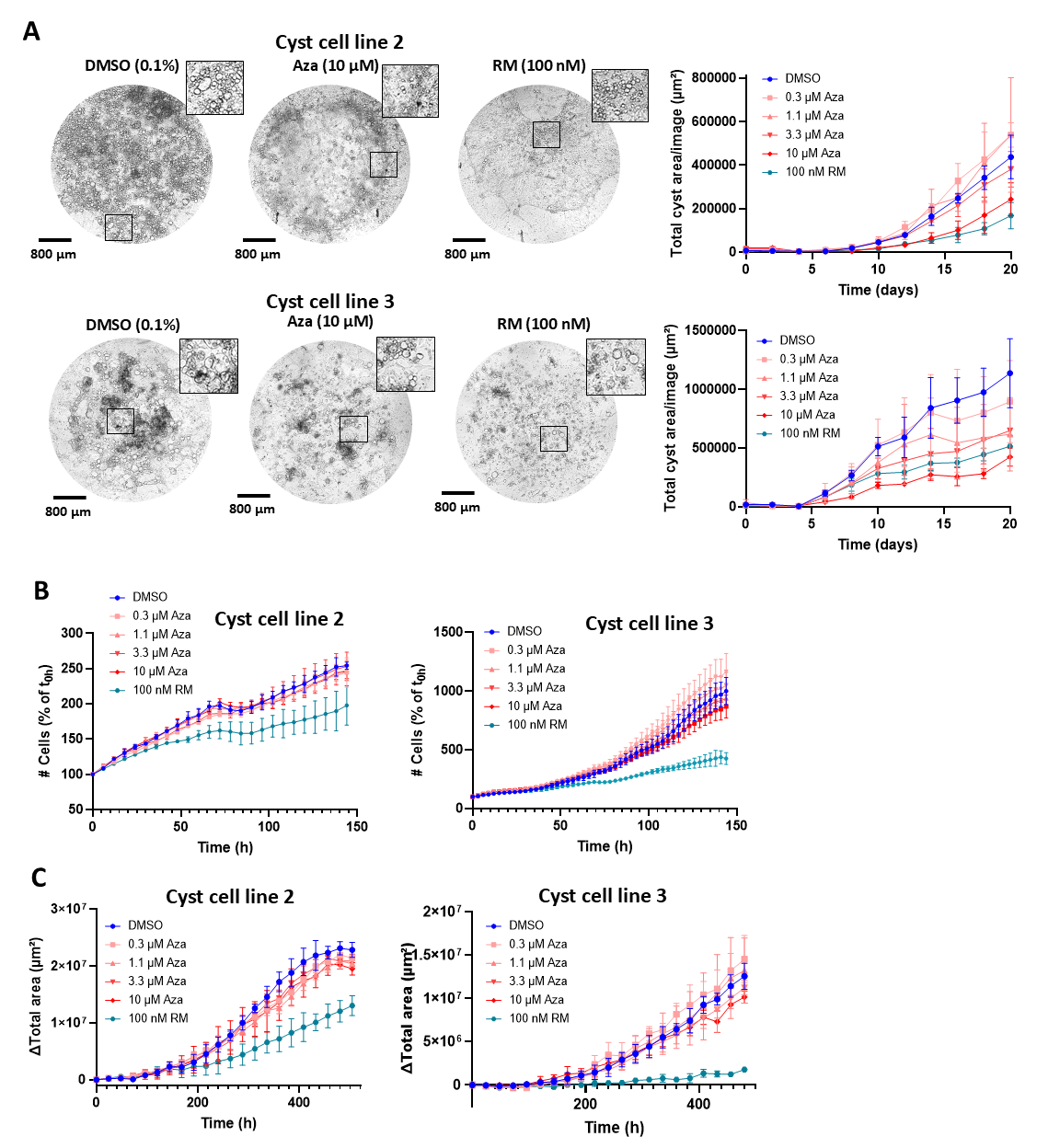
